# Behavioral and neuropathological features of Alzheimer’s disease are attenuated in 5xFAD mice treated with intranasal GHK peptide

**DOI:** 10.1101/2023.11.20.567908

**Authors:** Matthew Tucker, Gerald Yu Liao, Joo Young Park, Manuela Rosenfeld, Jackson Wezeman, Ruby Mangalindan, Dan Ratner, Martin Darvas, Warren Ladiges

## Abstract

Efforts to find disease modifying treatments for Alzheimer’s disease (AD) have met with limited success in part because the focus has been on testing drugs that target a specific pathogenic mechanism. Multiple pathways have been implicated in the pathogenesis of AD. Hence, the probability of more effective treatment for AD is likely increased by using an intervention that targets more than one pathway. The naturally occurring peptide GHK (glycyl-L-histidyl-L-lysine), as a GHK-Cu complex, supports angiogenesis, remodeling, and tissue repair, has anti-inflammatory and antioxidant properties, and has been shown to improve cognitive performance in aging mice. In order to test GHK-Cu as a neurotherapeutic for AD, male and female 5xFAD transgenic mice on the C57BL/6 background at 4 months of age were given 15 mg/kg GHK-Cu intranasally 3 times per week for 3 months until 7 months of age. Results showed that intranasal GHK-Cu treatment delayed cognitive impairment, reduced amyloid plaques, and lowered inflammation levels in the frontal cortex and hippocampus. These observations suggest additional studies are warranted to investigate the potential of GHK-Cu peptide as a promising treatment for AD.

## Introduction

Alzheimer’s disease (AD) is a complex disease and efforts to find effective treatments have met with limited success in part because the focus has been on testing drugs that target a specific pathogenic mechanism. The probability of effectively targeting several mechanistic pathways would be greatly increased by using a drug that individually targets more than one pathway (Ladiges, 2014; Rosenfeld and Ladiges, 2022).

GHK (glycyl-L-histidyl-L-lysine) is a naturally occurring peptide released from secreted protein acidic and rich in cysteine (SPARC) during proteolytic breakdown (Lane et al., 1994). In the event of an injury, GHK supports angiogenesis, remodeling, and tissue repair as a copper complex (GHK-Cu) (Pickart and Margolina, 2018; Badenhorst et al., 2016; Zhang et al., 2018). The peptide is clinically approved as a topical application for age-related skin conditions and promoted mainly as a skin rejuvenation drug (Dou et al., 2020). A previous search, using the Connectivity Map developed by the Broad Institute, found that GHK could alleviate impairment of TGFβ1 signaling (Pickart et al., 2017), a pathway associated with Aβ deposition and neurofibrillary tangle formation (Bosco et al., 2013). In addition, GHK has been shown to be an endogenous antioxidant by decreasing hydroxyl and peroxyl radicals (Sakuma et al., 2018), and improve cognitive performance in aging mice (Rosenfeld et al., 2023).

Targeted neurotherapeutics face a challenge in penetrating the blood brain barrier (BBB) (Ballabh et al., 2004; Hur et al., 2020; Uğurlu et al., 2011). This inherent restriction has significantly limited the availability of effective AD treatments, and has triggered investigation into methods or strategies to bypass the BBB. In this regard, the intranasal route is a promising delivery method to obtain greater efficacy in maintaining optimal drug concentrations in the brain compared to parenteral injections (Mittal et al., 2013; Chauhan and Chauhan, 2015; Crowe et al., 2017; Kumar et al., 2017). The intranasal system uses olfactory epithelium and its associated neural pathways to deliver drugs in therapeutically relevant concentrations.

The 5xFAD transgenic mouse line has been genetically engineered to model features of AD through the expression of human amyloid precursor protein (APP) and presenilin-1 transgenes, which are associated with early-onset AD (Brody et al., 2017). Mutations in these genes are linked to the regulation of amyloid-beta folding, a crucial component of AD pathogenesis. Transgenic 5xFAD mice have amyloid neuropathology similar to characteristic amyloid-plaque pathology in human patients (Brody et al., 2017; Younkin, 1998), and have been shown to develop impairment in cognitive behaviors that correspond to part of the dementia syndrome in AD.

The aim of this study was to determine if intranasal delivery of GHK-Cu as a neurotherapeutic would effectively delay the onset of cognitive impairment and neuropathology seen in 5xFAD transgenic mice. The advantages of intranasal administration, including reduced dosage loss, enhanced precision in drug delivery, and expedited brain access, were used to show that GHK-Cu was able to mitigate cognitive impairment and features of AD neuropathology.

## Methods

### Animals and experimental design

C57BL/6 mice with the transgenic 5xFAD genotype (JAX, Bar Harbor, Maine) of both sexes were used. The 5xFAD mouse line has five mutations: the Swedish (K670N/M671L), Florida (I716V), and London (V717I) mutations in APP, as well as the M146L and L286V mutations in PSEN1. Collectively, these mutations induce the formation of amyloid plaques. Mice were bred and genotyped following standard procedures from the Jackson Laboratory (Bar Harbor, ME, USA). Mice were group-housed, up to five per cage, in a specific pathogen free facility verified through viral and bacterial tests (IDEXX Bioanalytics). Nestlets (Ancare Corp, Bellmore, NY) were provided for physical and mental stimulation. Mice were monitored for health daily and cages were changed biweekly. All experimental procedures were approved by the University of Washington IACUC.

The experiment started when mice were 4 months of age and ended when they were 7 months of age for a treatment period of three months (12 weeks). Transgenic 5xFAD and wild type mice of both sexes were stratified across a peptide treatment cohort and a saline control cohort.

### GHK-Cu dose and delivery

GHK was used as a GHK-Cu complex (Active Peptide, Cambridge, MA) at a dose of 15 mg/kg body weight. Unpublished observations in our lab suggested that GHK-Cu complex was more effective than unbound GHK while unbound Cu had no effect in several in vitro and in vivo experiments. However, Cu ions can induce toxicity in mice if dosage exceeds levels of 35 mg/kg (Liu et al., 2020). Within the GHK-Cu complex, Cu made up 14% of the total molecular content, so a 15 mg/kg GHK-Cu dose had 2.1 mg/kg copper.

Drug administration occurred under 3% isoflurane anesthesia. Mice were then positioned at a 10-15° decline, and a hand grip was applied to the back, tail, and neck. This lowered the head in relation to the body, which maximized the surface area of the olfactory epithelium for efficient dosage uptake (Arber et al., 2017). The drug was then drawn up using a micropipette with a clean tip and applied carefully to the rim of the mouse nostril, one drop at a time. The timing of droplet placement was synchronized with natural inhalation to allow for drops to settle on the olfactory and respiratory epithelium within the nasal cavity (Mantzavinos and Alexiou, 2017). Droplets were alternated between nostrils with each breath until a 20⍰L volume was administered, typically requiring three to four droplets per mouse. Mice were then maintained in the declined position for an additional minute, during which stimulation was applied to the sternum region to enhance volume uptake. Intranasal GHK-Cu was administered three times per week using this procedure for the duration of the study.

### Behavioral Tests

Cognitive function was assessed by two behavioral tests, a spatial reference memory task (Y-Maze) and a spatial navigation learning task (Box Maze).

#### Y-Maze Task

The Y-Maze was performed as previously described (Darvas et al., 2009). Briefly, it consisted of three equally spaced arms at 120° angles from each other, each enclosed by raised walls. There was no escape option within the maze, and mice were allowed to freely explore it for five minutes while the path they took through the three arms was tracked by recording their entries into each arm. After completing the Y-maze, the mice were temporarily removed and placed in a separate resting cage while the maze was sterilized. All littermates underwent the assessment in a similar manner and were collectively returned to their home cage at the end of the trial. The Y-Maze test assessed spontaneous alternation, an indicator of working memory, as it measures the tendency of mice to explore novel arms of the maze rather than revisitations. Data from the trials administered at weeks four, eight, and twelve of the study were recorded. Spontaneous alternation data were expressed as a percentage calculated as the number of times a mouse completed a triad or loop through all three arms during the trial divided by the number of entries minus two (Miedel et al., 2017).

#### Box Maze Task

This task was performed as previously described (Darvas et al., 2020). Briefly, mice were introduced to a large foil-lined box with a bright overhead light, which served as a stressor. Each of the four walls of the box had two floor-level escape holes, with seven of them blocked. One hole was left open and connected to an S-curved escape tube leading to a darkened empty mouse cage. The escape hole setup was designed to create the illusion that each hole led into darkness so that mice had to rely on learning how to find the correct escape hole entrance to navigate an escape. Each mouse underwent three consecutive trials, each with a time limit of 180 seconds. Data were recorded from the trials, measuring the time in seconds it took each mouse to find and fully enter the escape hole.

##### Geropathology

At the conclusion of the twelve-week (3 months) treatment period, mice were euthanized using CO_2_ and tissues were collected and processed. Sections of the brain and other major organs were rapidly frozen in liquid nitrogen and stored at -80°C. The remaining brain sections, and systemic organs were fixed in 10% buffered formalin for 48 hours. Brain tissues were placed in PBS for 24 hours before being embedded in paraffin wax and sectioned onto histology slides. Brain section slides were stained with Congo red to evaluate amyloid plaques by averaging the number of plaques in ten different fields under 10x magnification in multiple areas of the brain blindly by two separate observers. Additional staining was performed using an Abcam immunohistochemistry kit for MCP-1 as a measure of general neuroinflammation. Stained slides were analyzed using Qu-Path digital imagining analysis to measure staining intensity as previously described (Lee et al., 2020).

### Data Analysis

Multiple behavioral groups were compared using a one-way analysis of variance (ANOVA) test. For the Box Maze, main trial-number effects were analyzed using the Bonferroni post-hoc multiple comparisons test to quantify the extent of latency differences. A two-tailed t-test was employed to compare results between cohorts for neuropathological data. All data were analyzed using GraphPad Prism software (version 10.0.3; GraphPad Software Inc., San Diego, CA, USA) as standard error of the mean with statistical significance at p< 0.05.

## Results

### Intranasal GHK-Cu peptide improved cognitive performance in transgenic 5xFAD mice

Both male and female transgenic mice treated with intranasal GHK-Cu had higher alternation percentages in the Y maze, indicating improved cognitive performance, beginning as early as the second month of treatment, and continuing through the third month when the study ended, compared to transgenic mice treated with intranasal saline (Figure 1A and B). Intranasal treatment with GHK-Cu was effective in preventing the cognitive impairment that was observed in saline treated transgenic mice and resulted in a cognitive performance level comparable to non-transgenic wildtype mice.

**Figure 1.**
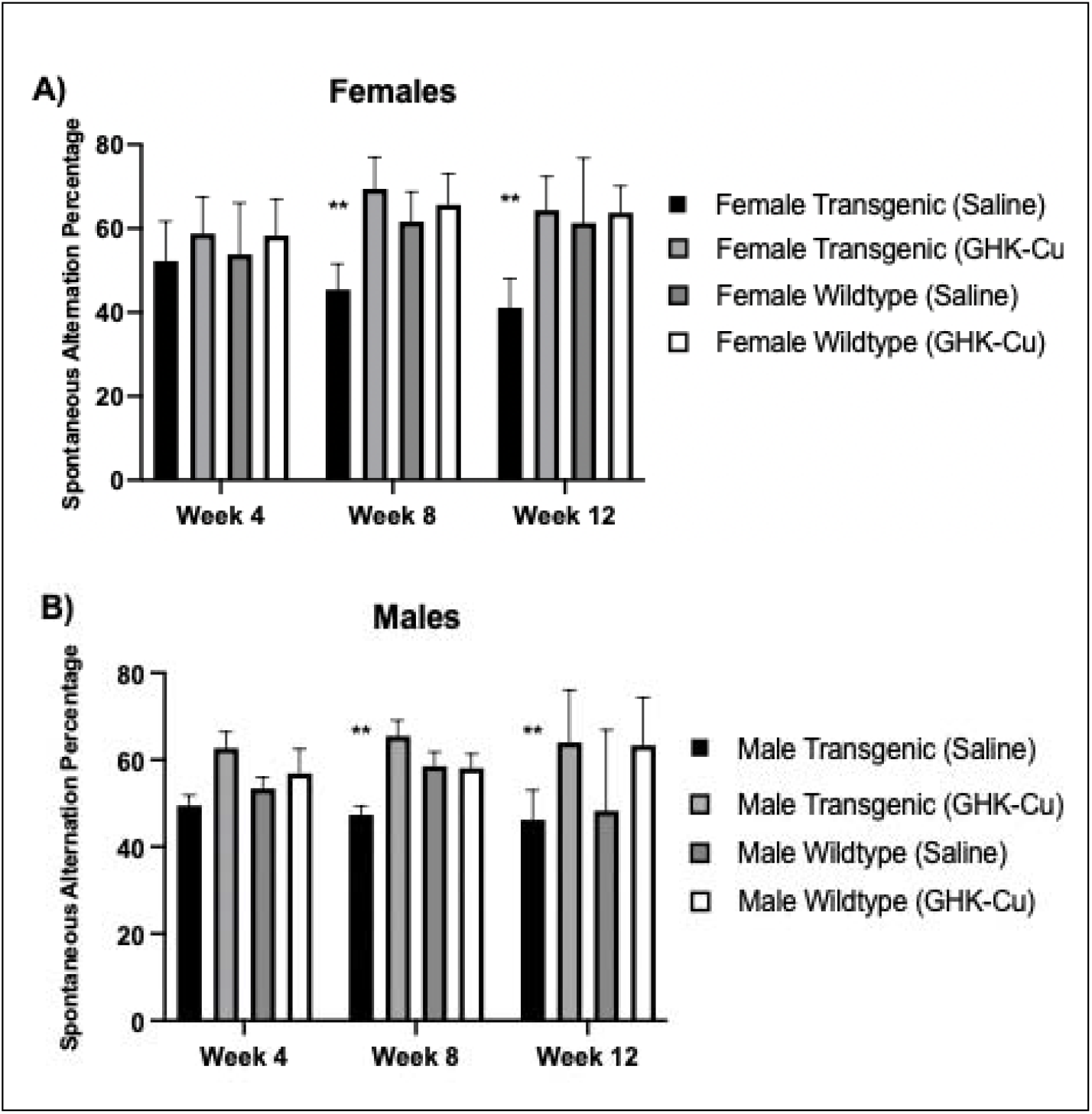
Y-maze percent alternation as a measure of cognitive function over 12 weeks of intranasal GHK-Cu treatment. Cognitive performance was evaluated relative to the 50% threshold. **A)** A higher alternation percentage was observed for female transgenic mice treated with intranasal GHK-Cu compared to transgenic females treated with intranasal saline at weeks 8 and 12. **B)** A similar pattern was observed for male transgenic mice treated with intranasal GHK-Cu compared to mice receiving intranasal saline at weeks 8 and 12. ** p≤0.01. N= 10-12/cohort. Transgenic = 5xFAD.

Box maze results after 12 weeks, when mice were 7 months of age, showed that transgenic mice treated with intranasal GHK-Cu had reduced escape latencies compared to transgenic mice treated with intranasal saline, reflecting decreased escape times over three trials and indicating improved learning capacity (Figure 2A and B). The effectiveness of intranasal GHK-Cu was seen equally in males and females and learning capacity was similar to that seen in non-transgenic (wild type) cohorts with or without intranasal GHK-Cu peptide.

**Figure 2.**
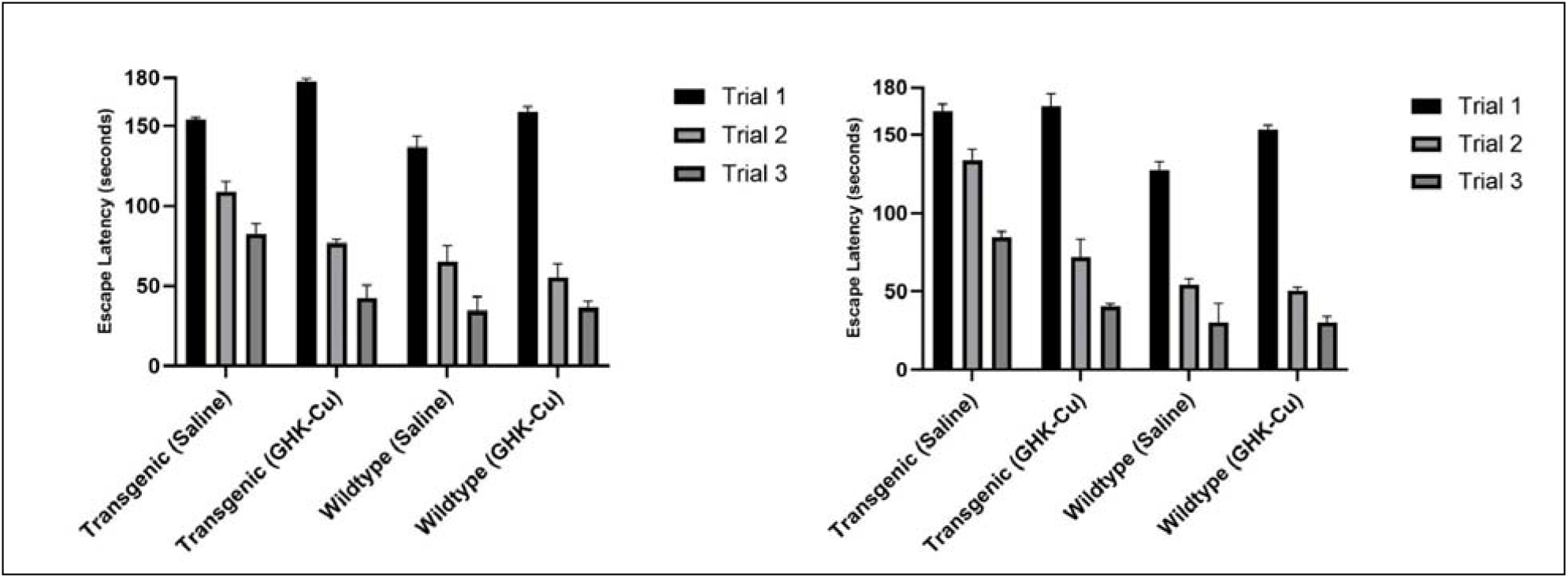
Transgenic 5xFAD mice treated with intranasal GHK-Cu for 12 weeks had faster escape times in the Box maze spatial navigation learning task. A decrease in escape latency, indicating faster escape time, was observed in female **(A)** and male **(B)** transgenic mice treated with intranasal GHK-Cu compared to transgenic mice given intranasal saline at trials 2 and 3, similar to both non-transgenic wild type cohorts. p≤0.01. N= 10-12/cohort. Transgenic= 5xFAD.

### Intranasal GHK-Cu peptide attenuated features of neuropathology in 5xFAD mice

The 5xFAD mouse genotype is characterized by the onset of amyloid plaques at 3 to 4 months of age, which progress to substantial and densely concentrated lesions in the brain (Oakley et al., 2006). Using a Congo Red stain, transgenic mice treated with intranasal GHK-Cu exhibited a significant reduction in amyloid plaques compared to transgenic mice treated with intranasal saline, irrespective of sex (Figure 3). Both male and female transgenic 5xFAD mice displayed the development of amyloid plaques in frontal cortex and hippocampus. Among the transgenic cohorts, those treated with intranasal GHK-Cu had significantly fewer detectable plaques in comparison to those receiving intranasal saline. Visual observation suggested a pattern where the plaques in saline-treated mice were generally larger and more densely stained compared to the plaques in GHK-Cu-treated cohorts (Figure 4). Wild-type (control) littermates did not display any amyloid plaques, consistent with their genotype.

**Figure 3.**
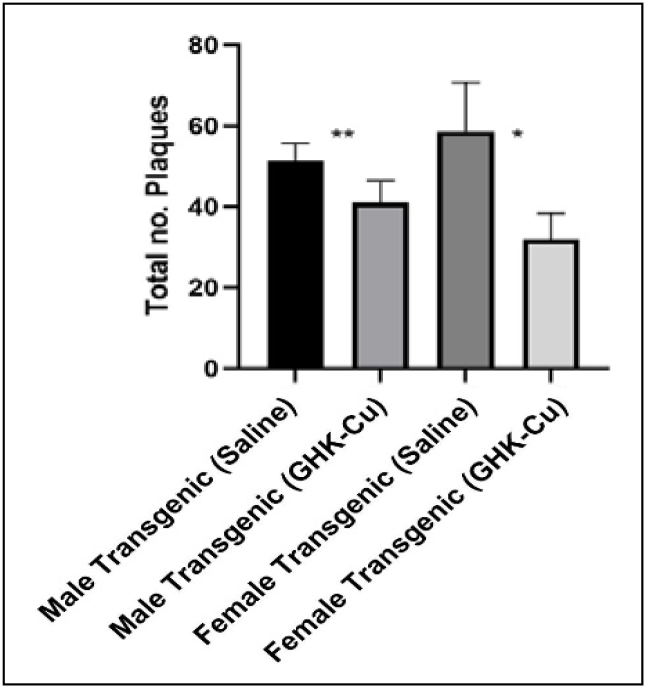
Amyloid plaques were reduced in the frontal cortex of 5xFAD mice following intranasal treatment with GHK-Cu. Quantitative visual count analysis of Congo Red-stained frontal cortex sections from both male and female mice showed a reduction in amyloid plaques in intranasal GHK-Cu-treated transgenic mice compared to intranasal saline-treated transgenic mice. *p≤0.15; **p≤0.05. N= 10-12/cohort. Transgenic = 5xFAD.

**Figure 4.**
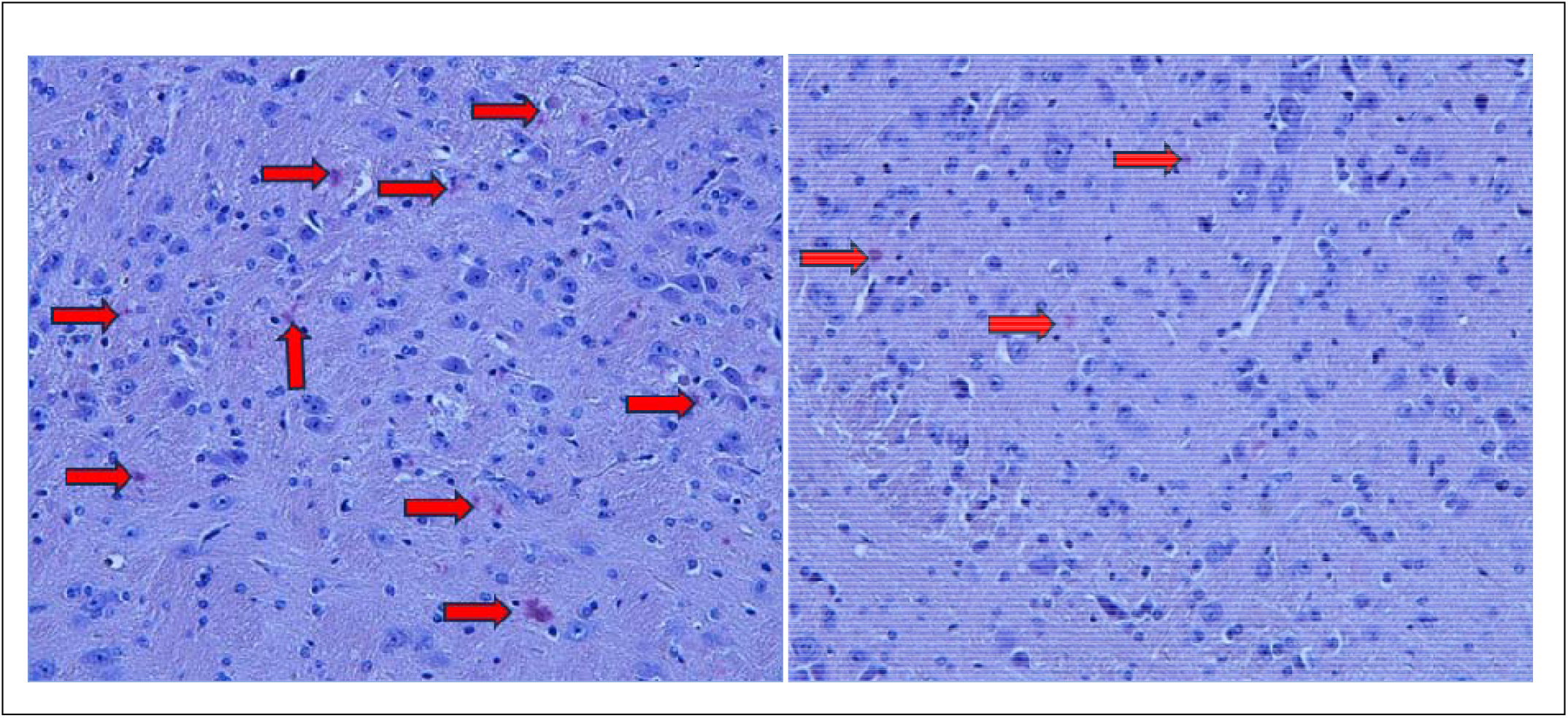
Representative Congo Red staining of frontal cortex sections from **A)** female transgenic mouse treated with intranasal saline or **B)** Female transgenic mouse treated with intranasal GHK-Cu. Transgenic = 5xFAD. Magnification 20X.

MCP-1 staining using IHC was conducted in the frontal cortex and the hippocampus. Results showed that both male and female transgenic 5xFAD mice that received intranasal GHK-Cu had decreased staining intensity for MCP-1 in both brain areas (Figure 5) indicating reduced neuroinflammation levels compared to transgenic mice treated with intranasal saline. Representative heat maps of frontal cortex visually show the respective staining intensity in female transgenic mice treated with intranasal saline or GHK-Cu (Figure 6A and B), compared to female wild type mice treated with intranasal saline or GHK-Cu (Figure 6C and D). It can be seen that visual staining intensity aligned with the quantitative values presented in Figures 4 and 5, and that 5xFAD mice treated with intranasal GHK-Cu had reduced levels of MCP-1 similar to wild type controls.

**Figure 5.**
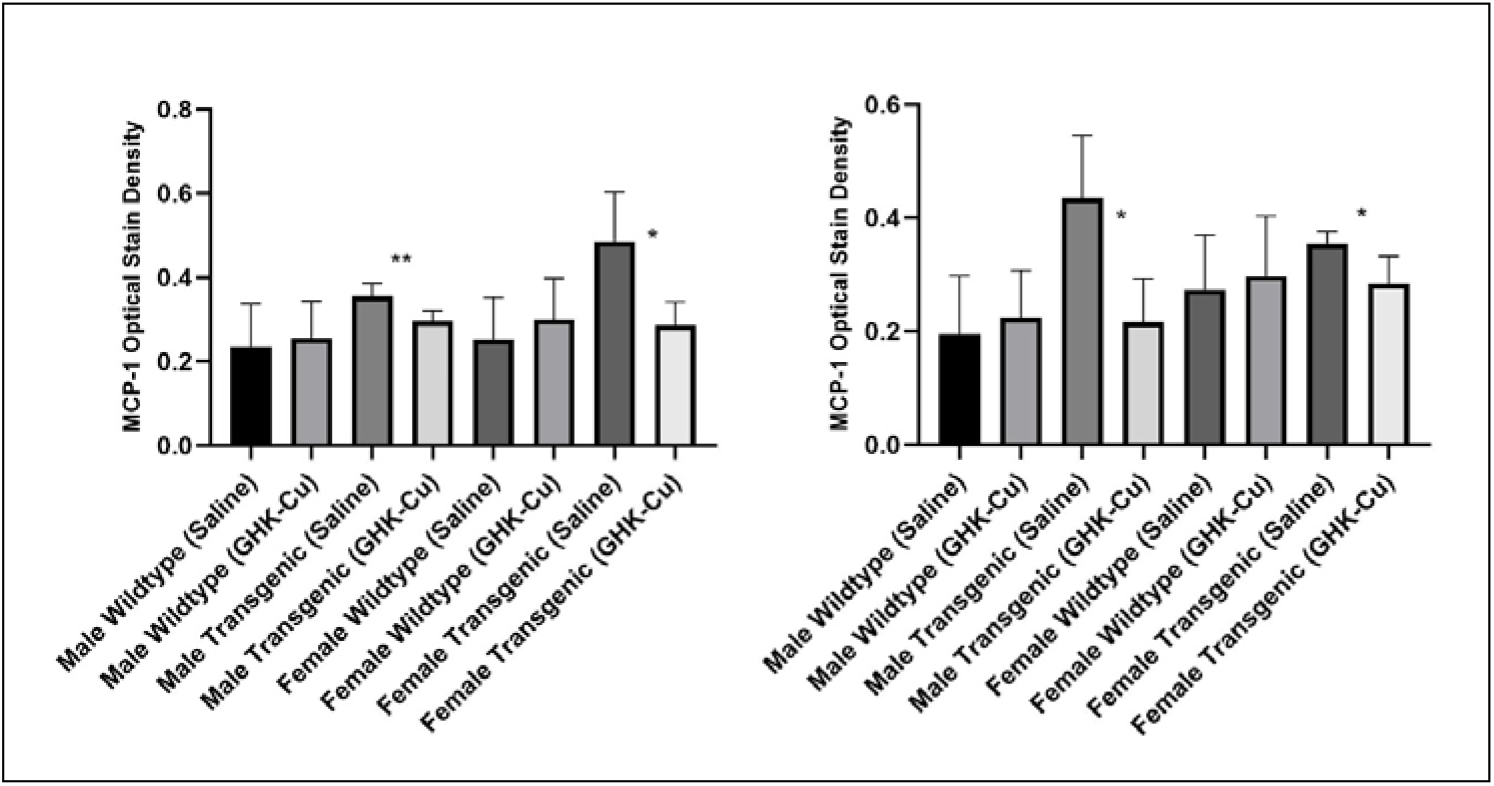
MCP-1 optical stain density in frontal cortex and hippocampus. **A**) Transgenic male and female mice treated with GHK-Cu displayed reduced optical stain density in the frontal lobe when compared to saline-treated cohorts. **B)** Transgenic male and female mice treated with GHK-Cu also exhibited lower optical stain density in the hippocampus in comparison to saline-treated counterparts. **p≤0.05, *p≤0.01. N= 6-10/cohort. Transgenic = 5xFAD.

**Figure 6.**
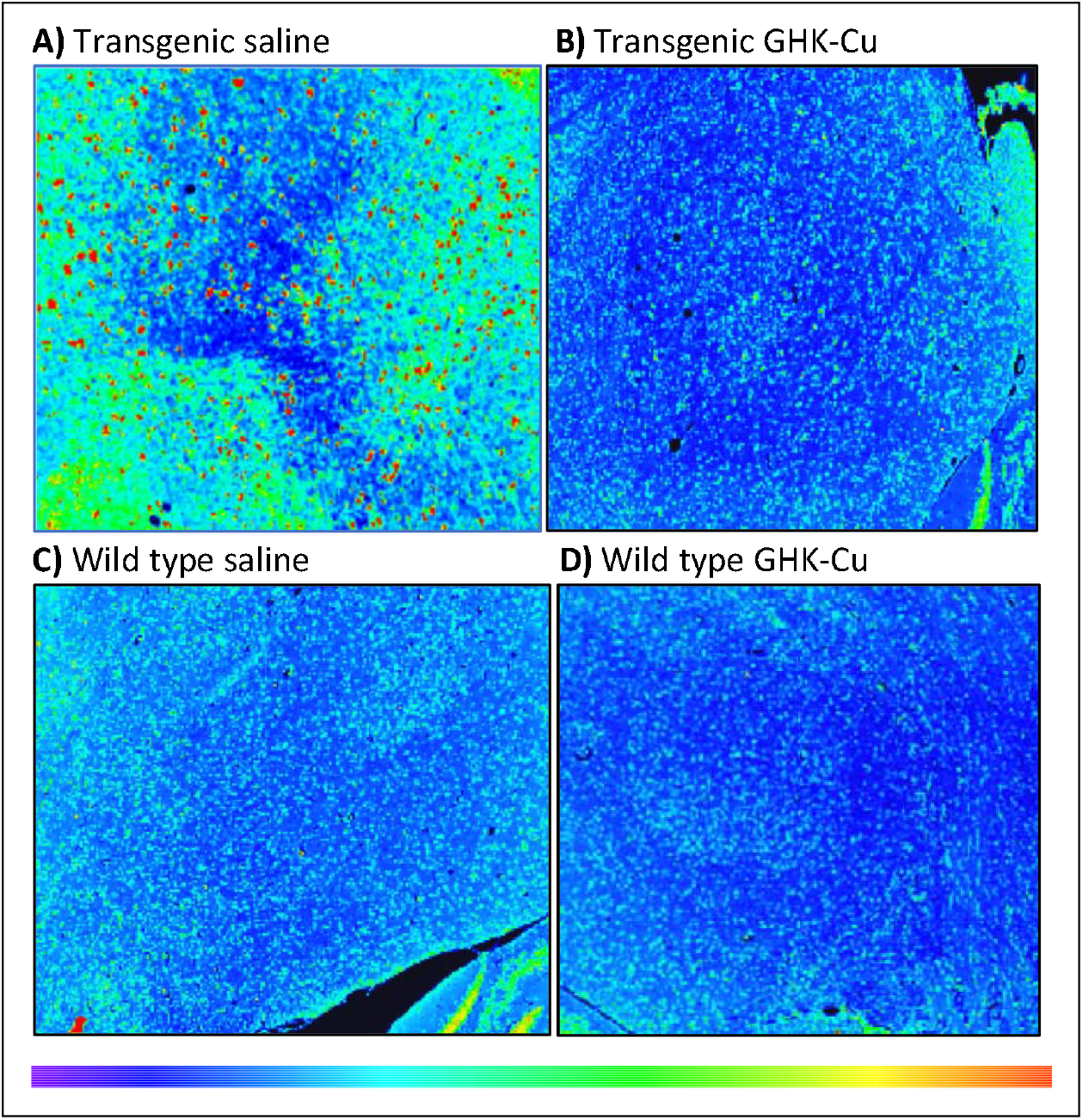
Q-Path generated heat map of an MCP-1 immunohistochemistry stain of frontal cortex from female mice after 12 weeks of treatment. **A**) Transgenic mouse treated with intranasal saline. **B**) Transgenic mouse treated with intranasal GHK-Cu. **C**) Wild type mouse treated with intranasal saline. **D**) Wild type mouse treated with intranasal GHK-Cu. The heat map legend indicates blue as low intensity staining all the way up to red as high intensity staining. Transgenic = 5xFAD; wild type = nontransgenic (age, strain, and sex matched controls).

## Discussion

This study showed that transgenic 5xFAD mice of both sexes exhibited improved cognitive performance after only 8 weeks of intranasal GHK-Cu treatment, compared to intranasal saline treated transgenic mice. This pattern continued through the 12-week treatment duration, corresponding to when the mice were 7 months old. Notably, cognitive improvement was coupled with a concurrent reduction in amyloid plaques and neuroinflammation in GHK-Cu-treated transgenic mice compared to saline-treated cohorts. Intranasal GHK-Cu treatment was therefore able to improve cognitive performance and reduce neuroinflammation levels to those of non-transgenic mice and significantly reduce the number of amyloid plaques.

Specifically, in the Y-maze spatial alternation test, GHK-Cu-treated transgenic mice of both sexes exhibited more spontaneous alternations, indicative of rescued working memory and prefrontal cortical functions (Kraeuter et al., 2018). Similar levels of spontaneous alternations were observed in both sexes of GHK-Cu-treated mice compared to other studies involving AD-related drug intervention (Cai et al., 2019; Jo et al., 2022; Morello et al., 2018). In the Box Maze task, GHK-Cu-treated 5xFAD mice of both sexes displayed decreased escape latencies, consistent with other reports on AD-related drug interventions (Mukherjee et al., 2019). However, evidence supporting the long-term effects of intranasal GHK-Cu on cognitive performance of 5xFAD mice is limited (Sosna et al., 2018; Morello et al., 2018), emphasizing the need for additional studies.

The accumulation of amyloid plaques is a pivotal factor associated with subsequent neuronal toxicity in AD pathogenesis, eventually leading to destruction of synaptic structures and severe cognitive deficits (Zhu et al., 2024). Our study revealed that GHK-Cu treated 5xFAD mice exhibited superior cognitive performance compared to saline-treated cohorts while also demonstrating a reduction in amyloid plaques in the frontal cortex and hippocampus. While the rescued cognitive abilities in GHK-Cu treated 5xFAD mice may be linked to diminished amyloid plaque formation, the extent of associated neurodegeneration in AD progression was not evaluated. Further investigations targeting this disparity are warranted based on a study protocol designed to assess neurodegeneration and neuronal loss specifically in 5xFAD mice approaching one year of age (Zhang et al., 2021).

Given that chemokine upregulation can result in chronic inflammation associated with onset and progression of age-related neurodegenerative diseases such as AD, our study employed MCP-1 staining in the frontal cortex and the hippocampus to assess the extent of possible inflammatory reactivity to amyloid plaques and microglial activity (Singh et al., 2021; Chen et al., 2016; Ishizuka et al., 1997). Our results demonstrated a notable decrease in MCP-1 staining intensity in both brain regions in 5xFAD mice that received intranasal GHK-Cu compared to saline-treated cohorts. The observed decrease in MCP-1 staining intensity suggests that intranasal administration of GHK-Cu in transgenic 5xFAD mice had a reducing effect on the AD-induced inflammatory phenotype and could serve as a reliable prototype to study drugs that slow or stop progression of AD (Sokolova et al., 2009).

The intranasal administration of GHK-Cu represents a novel approach aimed at overcoming the challenge of bypassing the blood-brain barrier, a common hurdle in traditional targeted neurotherapeutics (Ballabh et al., 2004; Hur et al., 2020; Uğurlu et al., 2011). By leveraging the olfactory epithelium and its associated neural pathways, this method capitalizes on the large surface area, efficient blood flow, and neural connections of the nasal mucosa to provide a relatively non-invasive approach towards efficient peptide delivery into the brain (Tulbah et al., 2023; Elkomy et al., 2023; Eissa et al., 2022; Elkomy et al., 2022; Abo El-Enin et al., 2022; Eid et al., 2019). The selection of a three-times-weekly administration schedule accounted for preventing adverse effects of daily anesthesia, and potential localized irritation and toxicity within the nasal mucosa (Tulbah et al., 2023; Elsenosy et al., 2020). While systemic toxicity of copper ions was carefully addressed, a comprehensive assessment of inflammation and inflammatory cell infiltration of the nasal epithelium in future studies would further validate the safety and non-toxicity of intranasal GHK-Cu administration.

The selection of the transgenic 5xFAD mouse model in our study was based on its ability to recapitulate an AD-like phenotype achieved through the expression of APP and PSEN1 transgenes (Brody et al., 2017; Youkin, 1998). Characterized by the onset of amyloid plaques at 3 to 4 months of age, and followed by cognitive impairment (Oakley et al., 2006), the 5xFAD mouse genotype validated these traits in our study. Among the transgenic cohorts, those treated intranasally with GHK-Cu exhibited improved cognitive performance and fewer, smaller, and less densely stained plaques compared to intranasal saline-treated cohorts. These observations provide the rationale to test intranasal GHK-Cu in an aging mouse model of AD, since AD is an age-related disease. An example is the induction of AD in aging mice using an AAV vector containing sequences of Aβ42 and mutant tau (Darvas et al., 2019).

In conclusion, this study demonstrates the positive therapeutic effect of intranasal GHK-Cu in transgenic 5xFAD mice, a widely used model for AD. Cognitive improvement was observed after 8 weeks of treatment, sustained through a 12-week study period, and accompanied by a concurrent reduction in amyloid plaques and neuroinflammation compared to intranasal saline-treated transgenic mice. Behavioral tests, including the Y-maze working memory task and the Box Maze spatial learning task, helped provide evidence that intranasal GHK-Cu treatment can enhance cognitive performance in a mouse model of AD. The intranasal delivery method allowed a non-invasive approach to efficient drug delivery. These findings suggest that intranasal GHK-Cu has the potential to attenuate features of AD, including cognitive decline, amyloid plaque accumulation, and neuroinflammation, and provides the rationale for further studies.

## Acknowledgements

This work was supported by National Institutes of Health grants R01 AG057381 (Ladiges and Darvas, co-PI’s) and R01 AG057381 (Ladiges PI).

## References

Abo El-Enin, Hadel A et al. “Lipid Nanocarriers Overlaid with Chitosan for Brain Delivery of Berberine via the Nasal Route.” Pharmaceuticals (Basel, Switzerland) vol. 15,3 281. 24 Feb. 2022, doi:10.3390/ph15030281

Arber, Charles et al. “Stem cell models of Alzheimer’s disease: progress and challenges.” Alzheimer’s research & therapy vol. 9,1 42. 13 Jun. 2017, doi:10.1186/s13195-017-0268-4

Badenhorst, Travis et al. “Physicochemical characterization of native glycyl-l-histidyl-l-lysine tripeptide for wound healing and anti-aging: a preformulation study for dermal delivery.” Pharmaceutical development and technology vol. 21,2 (2016): 152–60. doi:10.3109/10837450.2014.979944

Ballabh, Praveen et al. “The blood-brain barrier: an overview: structure, regulation, and clinical implications.” Neurobiology of disease vol. 16,1 (2004): 1–13. doi: 10.1016/j.nbd.2003.12.016

Bosco P, Raffaele Ferri, Maria Grazia Salluzzo, Sabrina Castellano, Maria Signorelli, Ferdinando Nicoletti, Santo Di Nuovo, Filippo Drago, Filippo Caraci. Role of the Transforming-Growth-Factor-β1 Gene in Late-Onset Alzheimer’s Disease: Implications for the Treatment.

Brody, David L et al. “Non-canonical soluble amyloid-beta aggregates and plaque buffering: controversies and future directions for target discovery in Alzheimer’s disease.” Alzheimer’s research & therapy vol. 9,1 62. 17 Aug. 2017, doi:10.1186/s13195-017-0293-3

Cai, Mudan et al. “Electroacupuncture attenuates cognition impairment via anti-neuroinflammation in an Alzheimer’s disease animal model.” Journal of neuroinflammation vol. 16,1 264. 13 Dec. 2019, doi:10.1186/s12974-019-1665-3

Chauhan, Mihir B, and Neelima B Chauhan. “Brain Uptake of Neurotherapeutics after Intranasal versus Intraperitoneal Delivery in Mice.” Journal of neurology and neurosurgery vol. 2,1 (2015): 009.

Chen, Wei-Wei et al. “Role of neuroinflammation in neurodegenerative diseases (Review).” Molecular medicine reports vol. 13,4 (2016): 3391–6. doi:10.3892/mmr.2016.4948

Crowe, Tyler P et al. “Mechanism of intranasal drug delivery directly to the brain.” Life sciences vol. 195 (2018): 44–52. doi: 10.1016/j.lfs.2017.12.025.

Darvas M, Keene D, Ladiges W. A geroscience mouse model for Alzheimer’s disease. Pathobiol Aging Age Relat Dis. 2019 May 14;9(1):1616994. doi: 10.1080/20010001.2019.1616994. PMID: 31143415; PMCID: PMC6522959.

Darvas, Martin et al. “A Novel One-Day Learning Procedure for Mice.” Current protocols in mouse biology vol. 10,1 (2020): e68. doi:10.1002/cpmo.68

Darvas, Martin et al. “Modulation of the Ca2+ conductance of nicotinic acetylcholine receptors by Lypd6.” European neuropsychopharmacology: the journal of the European College of Neuropsychopharmacology vol. 19,9 (2009): 670–81. doi: 10.1016/j.euroneuro.2009.03.007

Dou, Yan et al. “The potential of GHK as an anti-aging peptide.” Aging pathobiology and therapeutics vol. 2,1 (2020): 58–61. doi:10.31491/apt.2020.03.014

Eid, Hussein M et al. “Transfersomal nanovesicles for nose-to-brain delivery of ofloxacin for better management of bacterial meningitis: Formulation, optimization by Box-Behnken design, characterization and in vivo pharmacokinetic study.” Journal of Drug Delivery Science and Technology vol. 54, 101304. Dec. 2019, doi.org/10.1016/j.jddst.2019.101304

Eissa, Essam M et al. “Intranasal Delivery of Granisetron to the Brain via Nanostructured Cubosomes-Based In Situ Gel for Improved Management of Chemotherapy-Induced Emesis.” Pharmaceutics vol. 14,7 1374. 29 Jun. 2022, doi:10.3390/pharmaceutics14071374

Elsenosy, Fatma Mohamed et al. “Brain Targeting of Duloxetine HCL via Intranasal Delivery of Loaded Cubosomal Gel: In vitro Characterization, ex vivo Permeation, and in vivo Biodistribution Studies.” International journal of nanomedicine vol. 15 9517–9537. 30 Nov. 2020, doi:10.2147/IJN.S277352

Elkomy, Mohammed H et al. “Chitosan on the surface of nanoparticles for enhanced drug delivery: A comprehensive review.” Journal of controlled release : official journal of the Controlled Release Society vol. 351 (2022): 923–940. doi:10.1016/j.jconrel.2022.10.005

Elkomy, Mohammed H et al. “Intranasal Nanotransferosomal Gel for Quercetin Brain Targeting: I. Optimization, Characterization, Brain Localization, and Cytotoxic Studies.” Pharmaceuticsvol. 15,7 1805. 23 Jun. 2023, doi:10.3390/pharmaceutics15071805

Hur, Ga-Hee et al. “Effect of oligoarginine conjugation on the antiwrinkle activity and transdermal delivery of GHK peptide.” Journal of peptide science: an official publication of the European Peptide Society vol. 26,2 (2020): e3234. doi:10.1002/psc.3234

Ishizuka, K et al. “Identification of monocyte chemoattractant protein-1 in senile plaques and reactive microglia of Alzheimer’s disease.” Psychiatry and clinical neurosciences vol. 51,3 (1997): 135–8. doi:10.1111/j.1440-1819.1997.tb02375.x

Jo, Kyung Won et al. “Gossypetin ameliorates 5xFAD spatial learning and memory through enhanced phagocytosis against Aβ.” Alzheimer’s research & therapy vol. 14,1 158. 21 Oct. 2022, doi:10.1186/s13195-022-01096-3

Kraeuter, Ann-Katrin et al. “The Y-Maze for Assessment of Spatial Working and Reference Memory in Mice.” Methods in molecular biology (Clifton, N.J.) vol. 1916 (2019): 105–111. doi:10.1007/978-1-4939-8994-2_10

Kumar, Hitesh et al. “Intranasal Drug Delivery: A Non-Invasive Approach for the Better Delivery of Neurotherapeutics.” Pharmaceutical nanotechnology vol. 5,3 (2017): 203–214. doi:10.2174/2211738505666170515113936

Ladiges, Warren. “The quality control theory of aging.” Pathobiology of aging & age related diseases vol. 4 10.3402/pba.v4.24835. 23 May. 2014, doi:10.3402/pba.v4.24835

Lane, T F et al. “SPARC is a source of copper-binding peptides that stimulate angiogenesis.” The Journal of cell biology vol. 125,4 (1994): 929–43. doi:10.1083/jcb.125.4.929

Lee, Amanda et al. “QuPath. A new digital imaging tool for geropathology.” Aging pathobiology and therapeutics vol. 2,2 (2020): 114–116. doi:10.31491/apt.2020.06.024

Liu H, Guo H, Jian Z, Cui H, Fang J, Zuo Z, Deng J, Li Y, Wang X, Zhao L. Copper induces oxidative stress and apoptosis in the mouse liver. Oxidative Medicine and Cellular Longevity. 2020 Jan 11;2020.

Mantzavinos, Vasileios, and Athanasios Alexiou. “Biomarkers for Alzheimer’s Disease Diagnosis.” Current Alzheimer research vol. 14,11 (2017): 1149–1154. doi:10.2174/1567205014666170203125942

Miedel, Christian J et al. “Assessment of Spontaneous Alternation, Novel Object Recognition and Limb Clasping in Transgenic Mouse Models of Amyloid-β and Tau Neuropathology.” Journal of visualized experiments: JoVE,123 55523. 28 May. 2017, doi:10.3791/55523

Mittal, Deepti et al. “Insights into direct nose to brain delivery: current status and future perspective.” Drug delivery vol. 21,2 (2014): 75–86. doi:10.3109/10717544.2013.838713

Morello, Maria et al. “Vitamin D Improves Neurogenesis and Cognition in a Mouse Model of Alzheimer’s Disease.” Molecular neurobiology vol. 55,8 (2018): 6463–6479. doi:10.1007/s12035-017-0839-1

Morris, John C et al. “Assessment of Racial Disparities in Biomarkers for Alzheimer Disease.” JAMA neurology vol. 76,3 (2019): 264–273. doi:10.1001/jamaneurol.2018.4249

Mukherjee KK, Lee AY, Zhu L, Darvas M, Ladiges W. Sleep-deprived cognitive impairment in aging mice is alleviated by rapamycin. Aging Pathobiol Ther. 2019 Dec 30;1(1):5–9. doi: 10.31491/apt.2019.12.002. PMID: 35083443; PMCID: PMC8789090.

Oakley, Holly et al. “Intraneuronal beta-amyloid aggregates, neurodegeneration, and neuron loss in transgenic mice with five familial Alzheimer’s disease mutations: potential factors in amyloid plaque formation.” The Journal of neuroscience: the official journal of the Society for Neuroscience vol. 26,40 (2006): 10129–40. doi:10.1523/JNEUROSCI.1202-06.2006

Pickart, Loren, and Anna Margolina. “Regenerative and Protective Actions of the GHK-Cu Peptide in the Light of the New Gene Data.” International journal of molecular sciences vol. 19,7 1987. 7 Jul. 2018, doi:10.3390/ijms19071987

Pickart, Loren et al. “The Effect of the Human Peptide GHK on Gene Expression Relevant to Nervous System Function and Cognitive Decline.” Brain sciences vol. 7,2 20. 15 Feb. 2017, doi:10.3390/brainsci7020020

Rosenfeld, Manuela, and Warren Ladiges. “Pharmaceutical interventions to slow human aging. Are we ready for cocktails?.” Aging pathobiology and therapeutics vol. 4,2 (2022): 51–52. doi:10.31491/apt.2022.06.086

Rosenfeld, Manuela et al. “GHK peptide prevents sleep-deprived learning impairment in aging mice.” Aging pathobiology and therapeutics vol. 5,1 (2023): 33–35. doi:10.31491/apt.2023.03.109

Sakuma, Satoru et al. “The peptide glycyl--histidyl--lysine is an endogenous antioxidant in living organisms, possibly by diminishing hydroxyl and peroxyl radicals.” International journal of physiology, pathophysiology and pharmacology vol. 10,3 132–138. 25 Jun. 2018

Singh, Sanjiv et al. “MCP-1: Function, regulation, and involvement in disease.” International immunopharmacology vol. 101,Pt B (2021): 107598. doi:10.1016/j.intimp.2021.107598

Sokolova, Anna et al. “Monocyte chemoattractant protein-1 plays a dominant role in the chronic inflammation observed in Alzheimer’s disease.” Brain pathology (Zurich, Switzerland) vol. 19,3 (2009): 392–8. doi:10.1111/j.1750-3639.2008.00188.x

Sosna, Justyna et al. “Early long-term administration of the CSF1R inhibitor PLX3397 ablates microglia and reduces accumulation of intraneuronal amyloid, neuritic plaque deposition and pre-fibrillar oligomers in 5XFAD mouse model of Alzheimer’s disease.” Molecular neurodegeneration vol. 13,1 11. 1 Mar. 2018, doi:10.1186/s13024-018-0244-x

Tulbah, Alaa S et al. “Novel nasal niosomes loaded with lacosamide and coated with chitosan: A possible pathway to target the brain to control partial-onset seizures.” International journal of pharmaceutics: X vol. 6 100206. 12 Aug. 2023, doi:10.1016/j.ijpx.2023.100206

Uğurlu, Timuçin et al. “In vitro evaluation of compression-coated glycyl-L-histidyl-L-lysine-Cu(II) (GHK-Cu2+)-loaded microparticles for colonic drug delivery.” Drug development and industrial pharmacy vol. 37,11 (2011): 1282–9. doi:10.3109/03639045.2011.569934

Younkin, S G. “The role of A beta 42 in Alzheimer’s disease.” Journal of physiology, Paris vol. 92,3-4 (1998): 289–92. doi:10.1016/s0928-4257(98)80035-

Zhang, Heyu et al. “Glycine-Histidine-Lysine (GHK) Alleviates Neuronal Apoptosis Due to Intracerebral Hemorrhage via the miR-339-5p/VEGFA Pathway.” Frontiers in neuroscience vol. 12 644. 20 Sep. 2018, doi:10.3389/fnins.2018.0064

Zhang, Liansheng et al. “Assessment of neurodegeneration and neuronal loss in aged 5XFAD mice.” STAR protocols vol. 2,4 100915. 25 Oct. 2021, doi:10.1016/j.xpro.2021.100915

Zhu, Chenlu et al. “Rbm8a regulates neurogenesis and reduces Alzheimer’s disease-associated pathology in the dentate gyrus of 5×FAD mice.” Neural regeneration research vol. 19,4 (2024): 863–871. doi:10.4103/1673-5374.382254

